# Impact of Run-of-River Damming on Increasing Phytoplankton Biomass and Species Shift in a Large Amazonian River

**DOI:** 10.1101/2024.06.22.600178

**Authors:** Alfonso Pineda, Beatriz Melissa Campos, Felipe Zanon, Renata Felicio, Luzia C. Rodrigues

## Abstract

Run-of-river (ROR) dams, often perceived as having minimal environmental impact, can induce significant hydrodynamic changes that alter aquatic ecosystems. We investigated the impacts of an ROR dam on the Madeira River, the largest Amazon tributary, focusing on phytoplankton communities, their ecological implications, and related environmental factors. Our study examined changes in biomass and environmental factors (using General Linear Mixed Models - GLMM), species composition (using PERMANOVA) before and after damming, in both the main channel and tributaries (N=549 samples). We also identified indicator species associated with different damming phases and regions through an indicator value analysis. The results showed that, following dam construction, the phytoplankton community changed in both the main channel and tributaries, with a shift from lotic diatoms to lentic phytoflagellates. This transition was likely facilitated by altered hydrodynamics and possibly influenced by the decomposition of flooded vegetation in the dam’s influence zone. The decomposition of this vegetation could explain both the observed increase in oxygen consumption and the subsequent rise in phytoflagellate biomass after damming. However, despite the overall increase in phytoplankton biomass, the values remained within oligotrophic to mesotrophic conditions, consistent with the low nutrient concentrations recorded. However, we caution that the dam-created hydrodynamic conditions are optimal for phytoplankton growth, potentially exacerbating nutrient-related issues in the future. We recommend proactive management strategies to prevent nutrient enrichment from activities such as agriculture and livestock in isolated Amazon areas affected by dams, thereby mitigating potential water quality degradation linked to increased phytoplankton biomass.

## 1 Introduction

Hydroelectric energy is considered the most environmentally friendly option for meeting global energy demand. Hydroelectric projects have been constructed even in important ecological regions. For instance, in the Amazon region, the largest basin in the world, 140 dams are functioning or under construction, and 288 are planned (Latrubesse et al., 2017). Most of the hydroelectric projects built nowadays are run-of-river (ROR) structures, which in theory flood smaller areas than traditional storage reservoirs (Dias et al., 2018; Winemiller et al., 2016). However, even hydroelectric projects with a ROR dam can impact environmental conditions (Almeida et al., 2019; Moran, 2020; Tundisi, 2018), and biodiversity (Lees et al., 2016; Vasconcelos et al., 2022).

Run-of-river dams can have little effect on the main channel where they are built (Almeida et al., 2019; Zanon et al., 2024). However, they relate to the apparition of lentic or semi-lentic conditions in the back-flooded tributaries (Almeida et al., 2019). In some cases, the back-flooding area can correspond to 30% of the reservoir area (Almeida et al., 2019), and the total flooding area can be 64.5% larger than initially predicted by hydroelectric project including a ROR dam (Cochrane et al., 2017). These new lake-like conditions can harbor problematic species, such as mosquitoes which are vectors of diseases (Rodrigues et al., 2017), aquatic macrophytes that cover the water surface (Thomaz et al., 1999), and invasive species that exclude native ones (Lees et al., 2016).

The change from running water to standing water can also favor algae, as a longer water residence time enhances their reproduction. Standing water conditions together with high nutrient concentrations can trigger populational explosions of algae known as blooms (O’Neil et al., 2012). In turn, high algae biomass can threaten water quality (Ferrão-Filho and Kozlowsky-Suzuki, 2011), diversity (Krztoń et al., 2019), and human health (Pouria et al., 1998) since it decreases the transparency and oxygen of the water and even increases the concentration of toxins produced by some microalgae. Moreover, high algae biomass can degrade the services that we take advantage of from aquatic ecosystems such as recreation and fishing (Huisman et al., 2018).

The formation of standing waters in the Amazonian region could not trigger algae blooms *per se*, but together with the high temperature of the region, high nutrient input, and the hydrodynamics changes promoted by the dams may favor phytoplankton growth (Rodrigues et al., 2018; Thornton, 1990). For instance, human activities such as agriculture and cattle ranching in the Amazon increase phosphorous and nitrogen load in aquatic ecosystems (Tilman et al., 2002; Vale et al., 2019). This is particularly important given the increase in this type of enterprise in the Amazon in recent years (Carvalho et al., 2020; Gusso et al., 2017; Macedo et al., 2012; Vale et al., 2019).

Several studies have described the response of phytoplankton communities to damming in several regions worldwide (e.g., Okuku et al., 2016; Souza et al., 2016; Xiao et al., 2016; Znachor et al., 2020). However, it is no clear how the recently built and futures dams affects the spatial and temporal variation of phytoplankton communities in critical ecological regions such as the Amazon. Moreover, there exists a knowledge gap regarding the effect of damming on microalgae biomass of Amazonian Rivers. It is now crucial to determine whether the environmental changes induced by hydroelectric projects in the Amazon region are sufficient to promote the development of phytoplankton biomass and identify the favored species. This information is essential for alerting authorities to potential negative effects and for elucidating the appropriate management measures necessary to prevent or mitigate impacts associated with high phytoplankton biomass or unexpected species.

In this study, we assessed the influence of the Jirau dam on upstream biomass and community structure (species abundance) of phytoplankton in the Madeira River, one of the largest tributaries of the Amazon River. We collected data in the main channel and tributaries before and after the installation of the dam. Our objectives were to determine the effects of damming in phytoplankton biomass (structure and diversity) and identify which abiotic effects related to damming could explain eventual changes.

Since backwaters formed in tributaries after damming can exhibit lake-like conditions, we hypothesized that *i*) post-damming conditions in tributaries would favor an increase in phytoplankton biomass. Therefore, we expected an increase in biomass in tributaries, while no significant changes were anticipated in the main channel. Additionally, due to the shift in environmental conditions caused by damming, we hypothesized that *ii*) damming would alter the community structure (species abundance). Specifically, we anticipated differences in community structure before and after the damming, with species adapted to standing water conditions becoming representative after damming. Finally, we examined the significance of our findings in the context of environmental degradation resulting from human activities in the Amazon.

## 2 Methods

### 2.1 Study area

The Madeira River basin is in the Amazon region, between Peru, Bolivia, and Brazil, draining much of that region (Guyot et al., 1996). The Madeira River is among the ten rivers with the highest discharge in the world (Latrubesse et al., 2017)), it is the main tributary of the Amazon River and contributes significantly to the phosphorus load for the entire Amazon basin (Almeida et al., 2015). The Madeira River basin has one annual cycle with four well-defined hydrological periods, with well-established duration intervals between years: 1. The high-water period, which usually occurs from February to May; 2. The receding period, occurring from June to July; 3. low water, from August to November; and 4. rising period, occurring from December to January. This region has the largest tropical forest in the world, being responsible for much of the global biodiversity in addition to the importance of temperature and precipitation in South and Central America (Agudelo et al., 2019; Aragón, 2018).

The Madeira River has two dams installed, Santo Antônio and Jirau, but other dams are in the planning and approval phases (Latrubesse et al., 2017). The construction of the Jirau dam ended in July 2012, being considered a run-of-river, located in the city of Porto Velho, Rondônia state, Brazil. The Jirau reservoir has an annual average water residence time of approximately 3.0 days (Almeida et al., 2020). It is located 1,206 km from the mouth of the Madeira River and has an installed capacity of 3.750 MW, supplying more than ten million homes (Energia Sustentável do Brasil, 2020). In the study period, 2009 to 2017, the average flow was 19.409 m³/s, reaching a maximum average of 45.649 m³/s in the high-water period and a minimum average of 4.133 m³/s in the low-water period (ANA, 2022). For the study, we cover the area upstream of the Jirau hydroelectric plant, between the coordinates 9°67’73.05” S, 65°44’10.27” O, and 9°60’45.27” S, 64°91’75.55” O (Figure 1).

**Figure 1.**
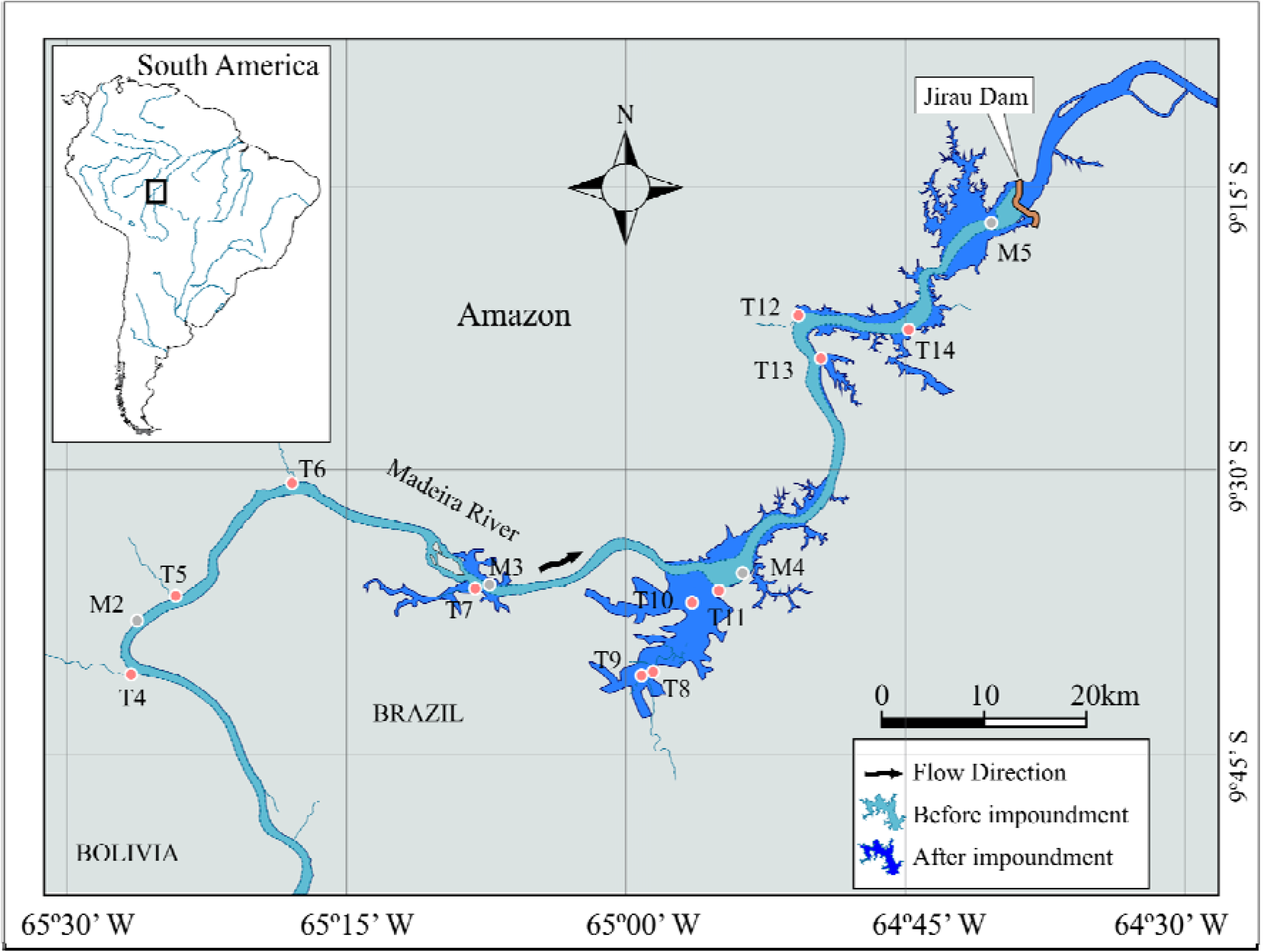
Map and location of sampling sites in the Jirau Hydroelectric Power Plant, in the Madeira River.

Run-of-river dams have been reported to have minimal influence on river flow and creating minor flooded regions (Pracheil et al., 2016; Souza et al., 2019). Nonetheless, the water level of the Madeira River has changed significantly since the Jirau dam was built. The water level ranged between 1 m and 20 m before damming, whereas it varied between 21 m and 29 m after damming. Also, the seasonal fluctuation in the Madeira River’s water level has decreased after damming (Supplementary material S1). The Madeira River’s rising water level has reduced flow velocity and induced stratification in its tributaries (Fearnside, 2013). The reservoir’s extension varies between 207 km^2^ during the dry season and 361.6 km^2^ during the wet season (Hartmann et al., 2023).

### 2.2 Data Samplings

Fifteen sampling stations were set up along the Madeira River (M2 to M5) and its tributaries (T4 to T14 – Figure 1). Water samples for phytoplankton and nutrient concentration assessments were collected directly using bottles at a depth of approximately 40 cm in the middle section of the Madeira Rivers and tributaries. At the same time, we measured chemical and physical variables that are critical to phytoplankton growth. Between 2009 and 2017, samples were taken during the high (March to May), rising (December to February), receding (June to August), and low (September to November) seasons. In 2010, 2011, 2016, and 2017, the campaigns were carried out quarterly, while from 2013 to 2015 they were carried out bimonthly. Five samplings were conducted in 2012. In total, we had 37 sampling campaigns (N = 550). Five samples were lost (M4 in campaign sampling 2, M3 in April 2010, T4 in April 2012, T10 in April 2013, and T5 in August 2014). The before-dam period is considered up until July 2012 (177 samples), while the after-dam period begins in October 2012 and continues through the remainder of the sampled periods (373 samples).

### 2.3 Phytoplankton

The sampled individuals were identified to the lowest possible taxonomic level, using specialized literature such as taxonomic articles, identification keys and books (Pineda et al., 2020a; van den Hoek et al., 1995; Komarek and Anagnostidis 1989, 1998, 2005). Phytoplankton samples were fixed with 1% acetic Lugol solution. Individuals (cell, coenobium, colonies, and filaments) were counted using an inverted microscope at 400X as described by Utermöhl (1958). Phytoplankton density was calculated according to APHA (American Public Health Association, 2012) and expressed as individuals per milliliter (ind.mL^-1^). Biomass (expressed in mm^3^.L-1) was obtained by multiplying the density of each taxon by its corresponding volume. The volume of individual cells was calculated using geometric models that were based on the shape of the cells (Hillebrand et al., 1999; Sun and Liu, 2003). The classification systems proposed by Reviers (2003) and Komárek & Anagnostidis (1998) were adopted for taxonomic identification at the level of class for eukaryotic algae and Cyanobacteria. The entire number of registered taxa was divided into five algal groups: green algae, cyanobacteria, diatoms, phytoflagellates, and xantophyceans. These groups are the most representative in freshwater ecosystems, accounting for most of the phytoplankton biomass. Additionally, they have shown sensitivity to environmental changes, making them well-suited for understanding how human activities affect biodiversity and water quality (Reynolds, 2006) (Reynolds, 2006). Although species from the Xanthophyceae class were identified, they were not included in the data analysis due to their low representativity, which made the analysis challenging.

### 2.4 Environmental data

We measured chemical and physical variables critical to the development of phytoplankton following the standard methods outlined in APHA (American Public Health Association, 2012).In situ measurements were conducted to assess water temperature (WT, °C), pH, inorganic carbon (IC, mg L^-1^), and alkalinity (Alk, mEq L^-1^). Water transparency was measured with a Secchi disk (m), and the euphotic zone (Zeu, m) was estimated as 2.7 times the Secchi depth (Cole, 1994). The maximum depth (Zmax, m) was also measured, and the Zeu:Zmax ratio was utilized as an indicator of light availability within the water column (Jensen et al., 1994).

Using collected water samples, we measured additional water characteristics at the sampled sites. Total iron concentrations (µg L^-1^) were determined following digestion using the phenanthroline method. Soluble reactive phosphorus (SRP, P-PO_4_^3-^, µg L^-1^), nitrate (N-NO_3_^-^, µg L^-1^), nitrite (N-NO_2_^-^, µg L^-1^), and ammonium (N-NH_4_^+^, µg L^-1^) concentrations were measured post-sample filtration. Phosphate concentrations were determined using the ascorbic acid method and spectrophotometric readings, while nitrate, nitrite, and ammonium concentrations were obtained through the cadmium reduction method, sulfanilamide method, and phenol method, respectively, for spectrophotometric measurements. Total dissolved inorganic nitrogen (DIN, µg L^-1^) was calculated as the sum of nitrite, nitrate, and ammonium concentrations. Reactive silica (µg L^-1^) concentrations were determined using the molybdate method. Biochemical oxygen demand (BOD, mg L^-1^) was assessed following sample incubation at a constant temperature (20 °C) and reading the dissolved oxygen concentrations after 5 days. Chemical oxygen demand (COD, mg L^-1^) determination involved the oxidation of organic matter using a boiling mixture of chromic acid and sulfuric acid (potassium bichromate in an acidic medium)(APHA, 2005).

### 2.5 Data analysis

We utilized Generalized Mixed Linear Models (GLMM) to investigate the effect of damming on the environmental variables in both the main channel and tributaries. To avoid issues with multiple pairwise tests, we included the sampling point as a random factor (Bolker et al., 2009). We selected an error distribution based on the better fit of the models, such as Gaussian, Beta, or Gamma.

To verify the effect of damming on the structure of the phytoplankton community, we utilized a Multivariate Permutation Analysis of Variance (PERMANOVA) on a matrix of species biomass (considering the Bray-Curtis distance). To determine whether damming effect interacted with hydrological periods, we considered the seasonality in the PERMANOVA. The analysis was performed separately for tributaries and the main channel. A Principal Coordinate Analysis (PCoA) was used to graphically depict the spatial and temporal ordinations. Additionally, to verify associations between species and damming phases (before and after) and regions (tributaries and main channel), we used the indicator value (IndVal) (Dufrêne and Legendre, 1997).We analyzed four groups that combined damming phases and regions. IndVal is a quantitative metric that accounts for both the relative abundance and frequency of a species within a specific condition and is not influenced by the abundance of other species. The significance of all indicator values was assessed using a Monte Carlo randomization process with 9999 permutations. A significant IndVal suggests that the species is more characteristic or prevalent in that condition. An IndVal value close to 1 indicates a strong association of the species with a particular condition (e.g., damming phase or region). In our study, this approach complemented the PERMANOVA analysis by allowing us to identify which species were more favored in each condition. For details on IndVal formula and its applications, see Borcard et al. (2018).

We utilized GLMM to assess the influence of damming on the biomass of the entire phytoplankton community and each of the five algal group in both the tributaries and the main channel. When we found that damming had a considerable influence on algal biomass, we examined into its link with seasonality. We also utilized GLMM to assess the influence of each standardized abiotic factor on algal biomass as well as on the presence-absence of each group. In this case, we modeled both biomass and the presence-absence of algal groups since several samples lacked a specific algae group, resulting in biomass values of zero (absence). We chose the Gamma family as the error distribution for modeling biomass and the zero-inflated distribution for occurrence data. To avoid issues with multiple testing, we incorporated the sampling point as a random effect in all models, as advised by Bolker et al. (Bolker et al., 2009). Visual examination validated the linearity assumption. The models’ findings were given in ANOVA style. To determine the trophic state before and after damming, the registered total biomass values were classified according to Vollenweider’s trophic classification (Vollenweider, 1968): oligotrophic (1 – 5 mm^3^L^-1^), mesotrophic (biomass = 3 – 5 mm^3^L^-1^), and eutrophic (10 mm^3^L^-1^).

Finally, we included thermal profiles from the main channel and tributaries to investigate thermal stratification following damming. These thermal profiles were limited to five sites (one in the Madeira River and four in tributaries) and were collected between 2012 and 2017. The statistical analyses together with their respective hypotheses were summarized in table 1. All analyzes were conducted utilizing the R program (R Development Core Team, 2024). Vegan package (Oksanen et al., 2007) was employed to perform PERMANOVA (adonis function) and to compute distance matrices (vegdist function). PCoA was performed with hnp package (Moral et al., 2017), GLMM (glmmTMB function) with glmmTMB package (Brooks et al., 2017) and plots with ggplot2 package (Wickham, 2016).

**Table 1.** Summary of data analyses and respective hypotheses.

| Hypotheses to test | Analyses used | References |
| --- | --- | --- |
| Hypothesis I: Changes in environmental characteristics and consequently changes in phytoplankton biomass | GLMM | Brooks, Mollie et al., 2017 |
| Hypothesis II: Change in phytoplankton composition between the pre- and post-dam phases | Indval | Dufrêne and Legendre, 1997 |
|  | PERMANOVA | Anderson, 2017 |

## 3 RESULTS

### 3.1 Environmental conditions

Both the main channel and tributaries exhibited changes in the environment after damming, but changes were more pronounced in the tributaries (Table S1, Figures 2 and 3). An increase in chemical oxygen demand (COD) was observed in both environments. The main channel showed a reduction in alkalinity, inorganic carbon, silica, conductivity, and pH, whereas tributaries displayed a decrease in light availability (Zeu and Zeu:Zmax) and iron, but an increase in depth, conductivity, inorganic carbon, temperature, and alkalinity. The variation of most environmental factors seemed to increase with damming at both the main channel and tributaries (Table 2), except for orthophosphate, BOD, and COD which decreased.

**Figure 2.**
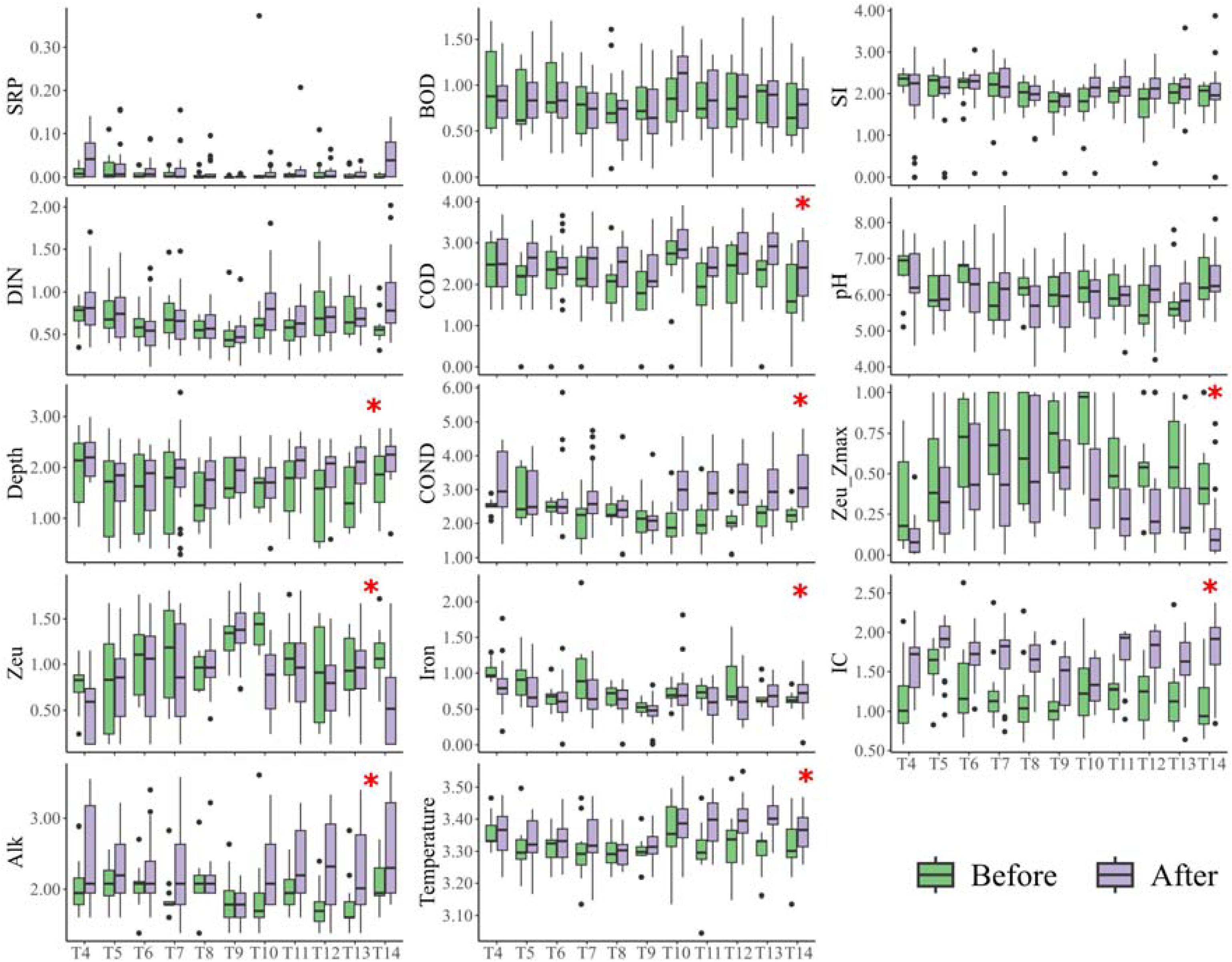
The effect of damming on the variation of environmental variables measured at the tributaries of the Madeira River. Asterisks (*) in the figure indicate a significant difference. The vertical axis displays the log-transformed values for each variable, except for pH and the ratio of Zeu:Zmax. *IC* – dissolved inorganic carbon; *BOD* – biochemical oxygen demand; *COD* – chemical oxygen demand; *DIN* – dissolved inorganic nitrogen; *Z_eu_:Z_max_* – light availability in the water column; *COND* – Conductivity; *Alk* – alkalinity; *Zeu* – euphotic zone; *SRP* – soluble reactive phosphorous; *SI* - Silica.

**Figure 3.**
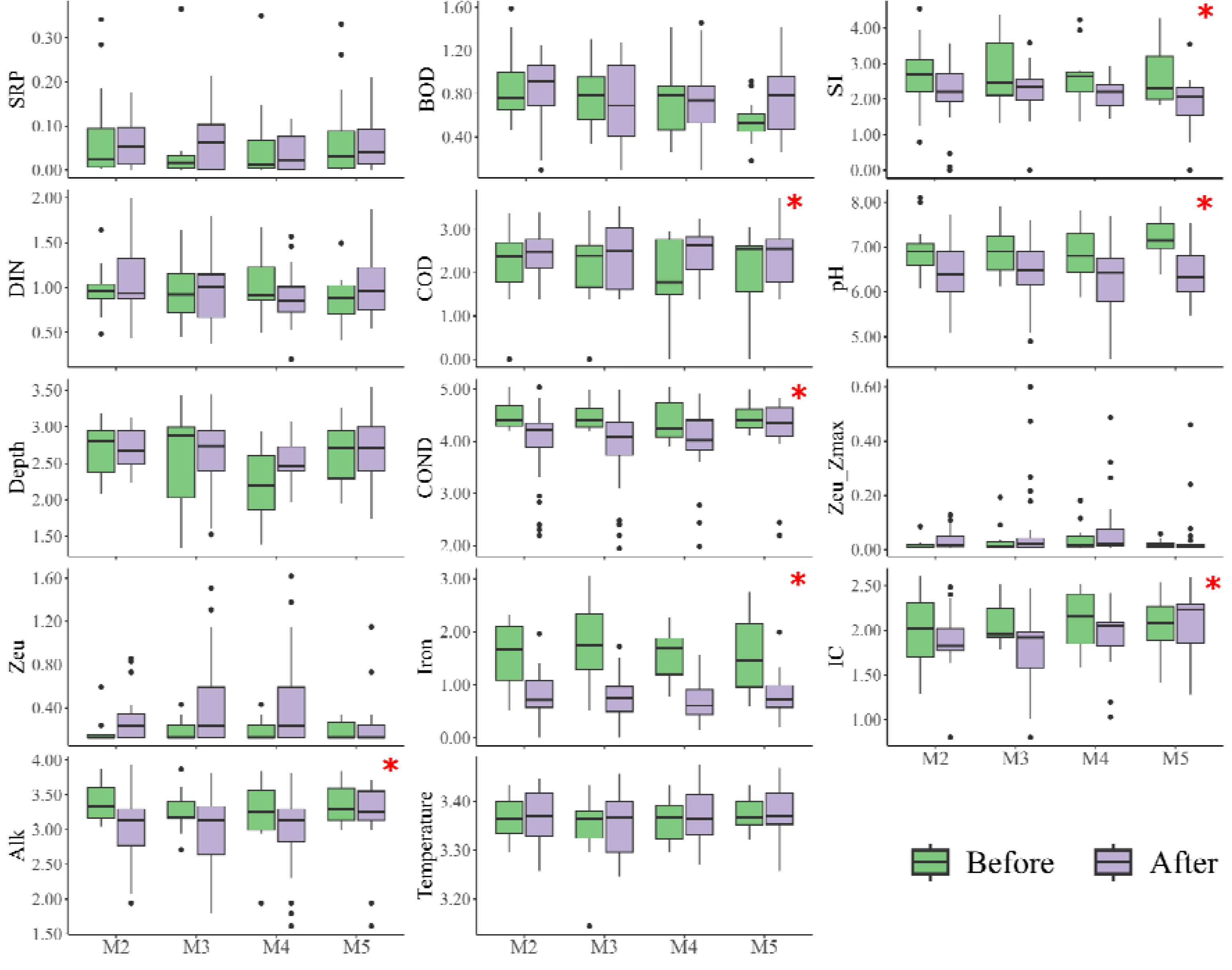
The effect of damming on the variation of environmental variables measured at Madeira River. Asterisks (*) in the figure indicate a significant effect of damming. The vertical axis displays the log-transformed values for each variable, except for pH and the ratio of Zeu:Zmax. The boxes in the plot represent the first and third quartiles, the horizontal bar represents the median, and the vertical bars indicate the minimum and maximum values, excluding any outliers, which are represented as individual points. IC – dissolved inorganic carbon; BOD – biochemical oxygen demand; COD – chemical oxygen demand; DIN – dissolved inorganic nitrogen; Zeu:Zmax – light availability in the water column; COND – Conductivity; Alk – alkalinity; Zeu – euphotic zone; SRP – soluble reactive phosphorous; SI - Silica.

**Table 2:** Mean and coefficient of variation (CV, %) of abiotic variables in the main channel and tributaries during the construction phases of the dam.

|  | Madeira |  | Tributaries |  |
| --- | --- | --- | --- | --- |
|  | Before | After | Before | After |
| Temperature (°C) | 27.91<br>(5.68) | 28.04<br>(6) | 26.61<br>(7.98) | 27.76<br>(7.32) |
| pH | 7<br>(7.7) | 6.41<br>(10.88) | 6.18<br>(11.84) | 6.06<br>(13.06) |
| Conductivity ($\mu\text{S.cm}^{-1}$) | 88.9<br>(32.17) | 66.35<br>(52.98) | 10.31<br>(75.87) | 25.54<br>(123.92) |
| $Z_{\text{eu}}:Z_{\text{max}}$ (m) | 0.03<br>(141.07) | 0.06<br>(184.55) | 0.59<br>(51.73) | 0.36<br>(85.52) |
| Euphotic zone - $Z_{\text{eu}}$ (m) | 0.22<br>(66.38) | 0.57<br>(136.02) | 2.09<br>(59.31) | 1.63<br>(73.58) |
| Orthophosphate ($\mu\text{g L}^{-1}$) | 0.09<br>(162.19) | 0.07<br>(85.63) | 0.01<br>(370.74) | 0.02<br>(159.5) |
| Dissolved inorganic nitrogen ($\mu\text{g L}^{-1}$) | 1.74<br>(52.95) | 1.89<br>(62.93) | 0.97<br>(61.4) | 1.13<br>(73.73) |
| Biochemical oxygen demand ($\text{mg L}^{-1}$) | 1.18<br>(66.06) | 1.28<br>(56.22) | 1.45<br>(66.4) | 1.39<br>(61.85) |
| Chemical oxygen demand ($\text{mg L}^{-1}$) | 10.11<br>(74.71) | 12.69<br>(63.76) | 9.47<br>(71.57) | 14.48<br>(65.46) |
| Iron ($\mu\text{g L}^{-1}$) | 5.05<br>(80.42) | 1.39<br>(80.54) | 1.25<br>(71.22) | 1.03<br>(61.62) |
| Alkalinity ($\text{mEqL}^{-1}$) | 28.22<br>(34.25) | 23.19<br>(45.47) | 6.71<br>(58.59) | 10.63<br>(72.73) |
| Inorganic carbon ($\text{mg L}^{-1}$) | 7.31<br>(36.15) | 6.31<br>(37.72) | 2.76<br>(71.73) | 4.53<br>(38.89) |
| Silica ($\mu\text{g L}^{-1}$) | 22.55<br>(107.92) | 9.28<br>(73.69) | 6.94<br>(43.68) | 7.94<br>(57.16) |

WT - water temperature - °C; pH; IC – dissolved inorganic carbon; BOD - biochemical oxygen demand; COD - chemical oxygen demand; DIN – dissolved inorganic nitrogen; $Z_{\text{eu}}:Z_{\text{max}}$ – light availability in the water column, SRP – soluble reactive phosphorus, Si -Silica.

### 3.2 Phytoplankton structure and biomass

In this study, we identified a total of 525 taxa, with the highest number of species belonging to green algae (236 species, mean of 0.095 ind.mL^-1^), Phytoflagellates comprised 118 species (mean of 0.145 ind.mL^-1^), followed by diatoms (86 species, mean of 0.057 ind.mL^-1^). Cyanobacteria accounted for 72 species (mean of 0.017 ind.mL^-1^), and xantophyceans were represented by 13 species (mean of 0.0004 ind.mL^-1^).

Before and after damming, the main channel recorded 119 and 167 species, respectively, with diatoms (mainly bacillariophyceans) exhibiting the highest richness both before (30) and after (37) damming. In the tributaries, the number of species increased from 307 before damming to 457 after damming, with zygnemaphyceans having the greatest richness both before (62) and after (94) damming.

PERMANOVA revealed a significant effect of damming on the phytoplankton diversity for the main channel (F = 1.87, p = 0.01) and tributaries (F = 12.89, p <0.01). In addition to impoundment, we also found a significant effect of hydrological periods on community diversity (for main channel, F = 1.22, p = 0.001; for tributaries, F = 2.09, p <0.01). Both the PERMANOVA results and their graphical representation (Figure 4) showed that damming altered communities and increased the spatial and temporal variation in species biomass in both the main channel and tributaries across all hydrological periods.

**Figure 4:**
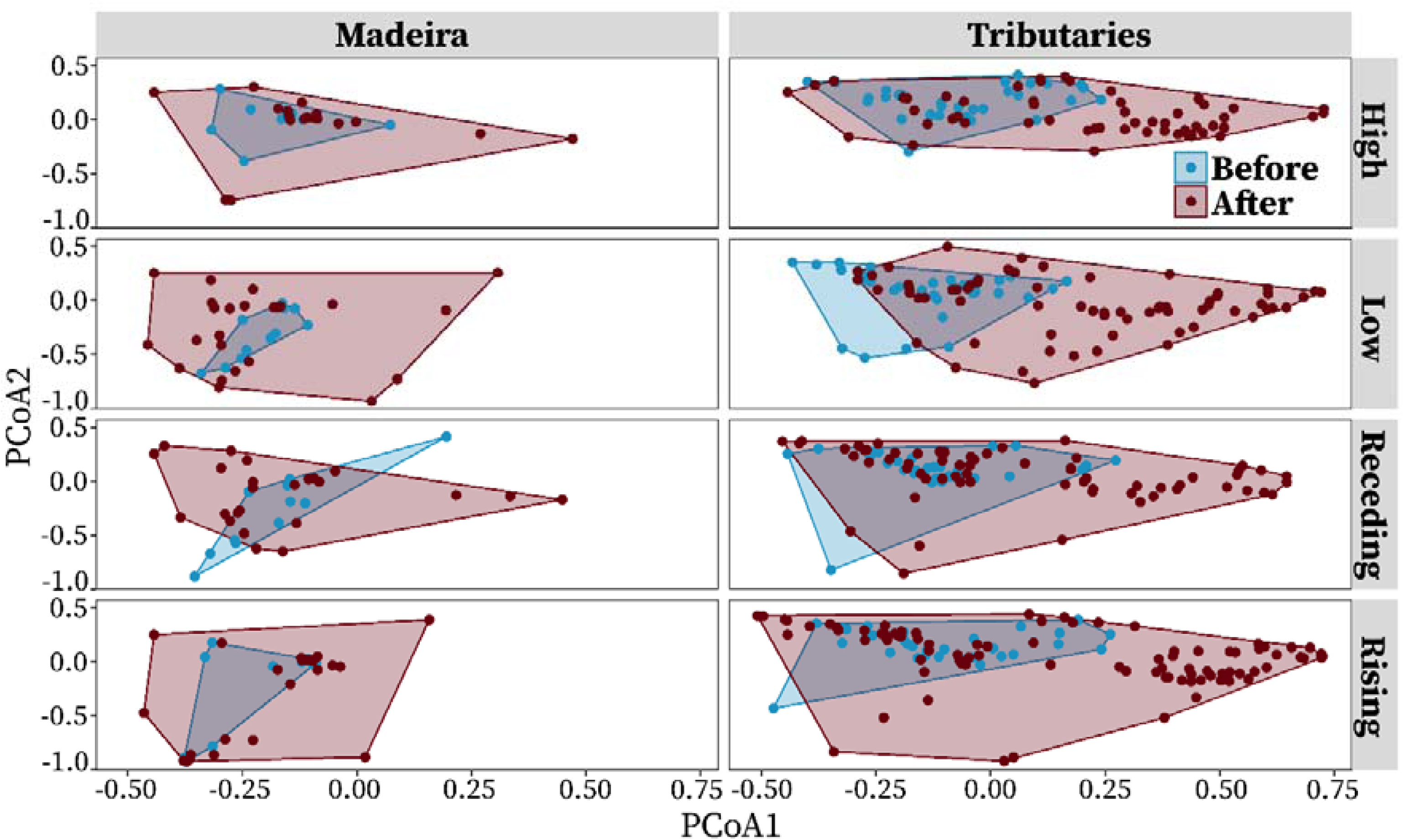
Temporal variation of phytoplankton community (species abundance) related to damming phases and hydrological periods at both the main channel and tributaries. In all cases, the community’s heterogeneity was higher in the after damming phase. The higher variation seemed to be related to the low water period.

IndVal analysis indicated that the number of species linked with tributaries increased after damming, while indicators of the Madeira River decreased. Diatoms, which are related to flowing waters, were found to be exclusive indicators of pre-damming conditions for both tributaries and the main channel. It is noteworthy that most of the indicator species associated with tributaries in the post-damming phase were indicative of standing water, specifically phytoflagellates (Table 3).

**Table 3.**
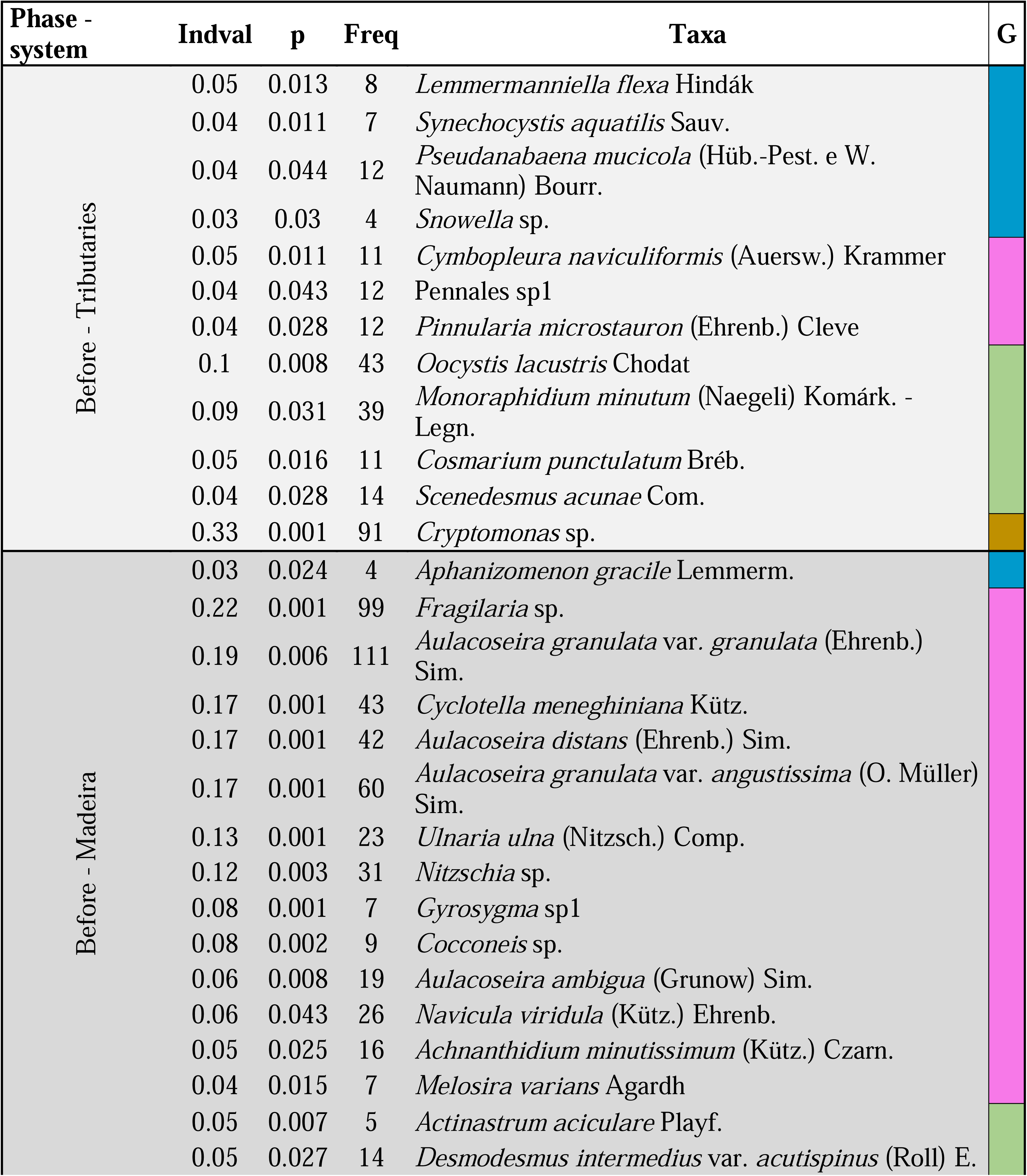
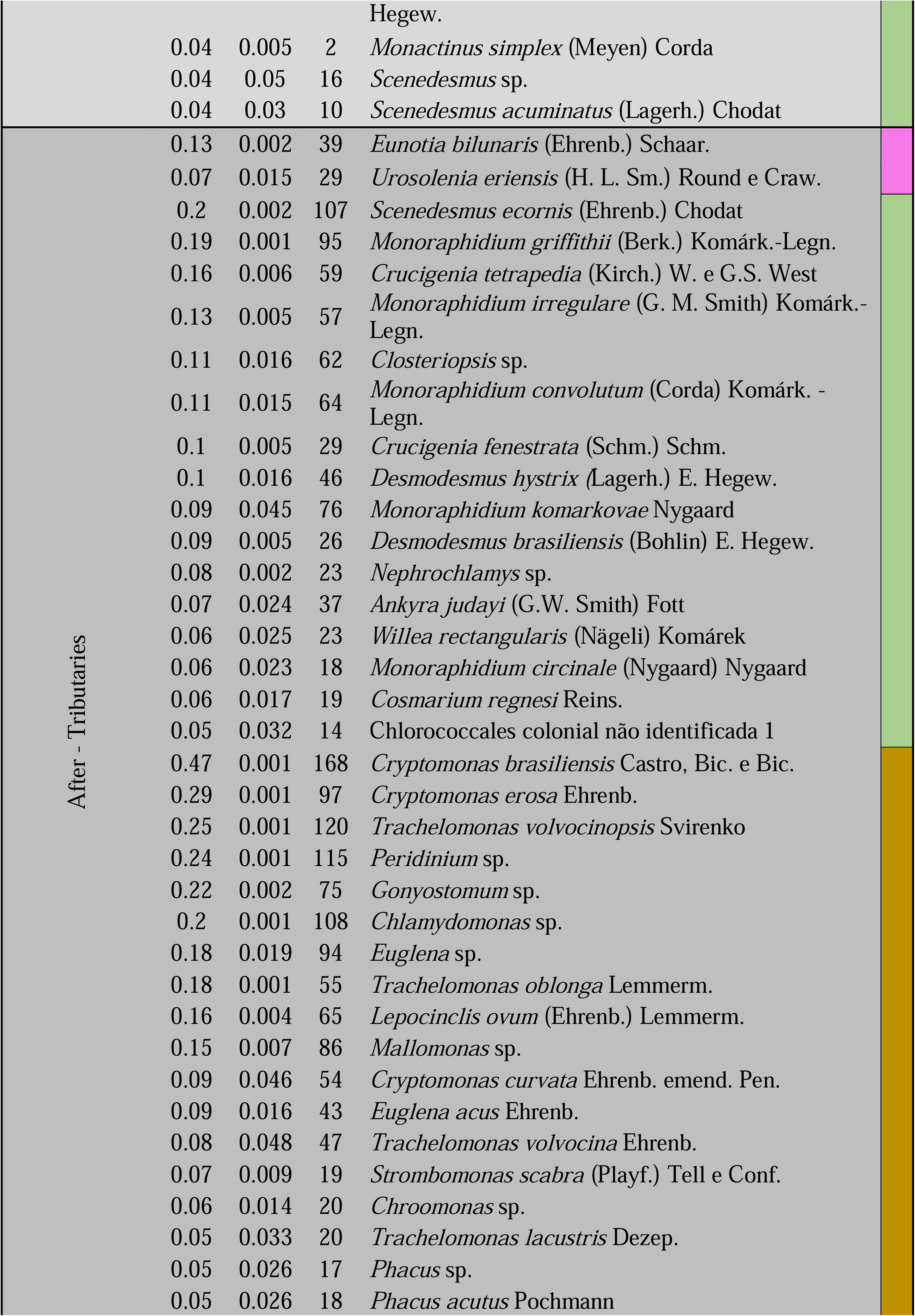
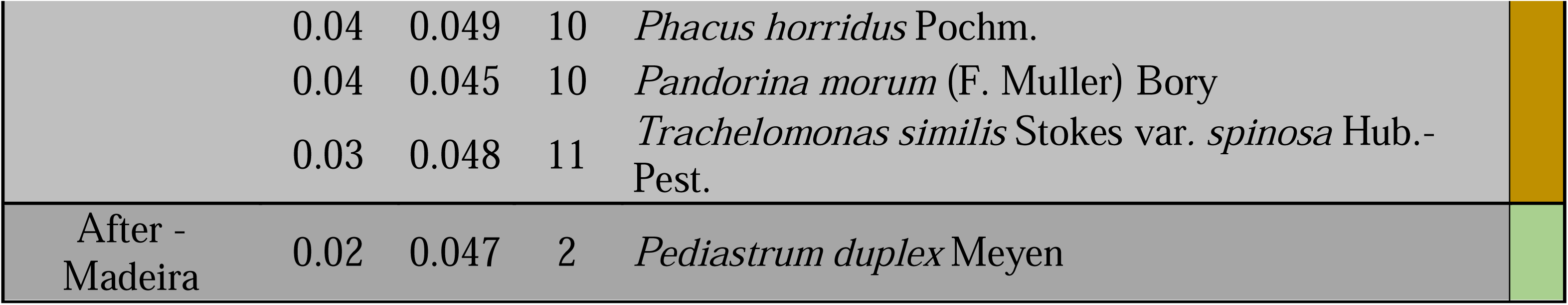
Species as indicators of the conditions at each damming phase (before and after) and system (tributaries and Madeira River). *IndVal*, indicator value based on the relative frequency and relative average abundance in the clusters Phase – system. The table shows only those species with a high probability of being an indicator (p < 0.05). *Freq*, specie **frequency** at each cluster. The taxonomic groups were colored differently to highlight the representativity of each group at each phase – system: blue – Cyanobacteria; pink – Diatoms; green – Green algae; golden – Phytoflagellates.

In the tributaries, the total biomass ranged from 0.0001 to 4.74 mm^3^L^-1^ before damming. After damming values varied between 0.0001 and 15 mm^3^L^-1^. Most samples (98% before and 96% after damming) showed a biomass lower than 2 mm^3^L^-1^. After damming, we observed biomass higher than 2 mm^3^L^-1^ (four samples), 3 mm^3^L^-1^ (four samples), 5 mm^3^L^-1^ (1 sample), and 10 mm^3^L^-1^ (one sample). In the Madeira River, the total biomass ranged between 0.001 and 3.38 mm^3^L^-1^ before damming, and after damming, values varied between 0.0001 and 0.97 mm^3^L^-1^. Biomass was less than 2 mm^3^L^-1^ in 98% of the samples before damming and 100% after damming. According to Vollenweider, the registered biomass conditions mainly ranged from oligotrophic (biomass = 1 – 5 mm^3^L^-1^) to mesotrophic (biomass = 3 – 5 mm^3^L^-1^).

GLMM revealed differences in total biomass between damming phases in both the main channel and tributaries (Figure 5A and Table S2). In the tributaries, total and phytoflagellates biomass increased (Figure 5A), with variations between phases depending on the season considered. Specifically, no differences were observed during the rising season for total biomass and at high water for phytoflagellates, while cyanobacteria showed differences only in the high and receding water seasons (Figure 5B and Table S3). In the main channel, total biomass decreased, along with diatoms and green algae biomass, while phytoflagellates increased (Figure 5A). Differences between the phases were consistent across season for phytoflagellates. In contrast, differences between the phases at the low- and rising-water seasons were observed for the total biomass and diatoms, at the rising- and high-water seasons for greens, and at the rising waters for cyanobacteria (Figure 5B).

**Figure 5:**
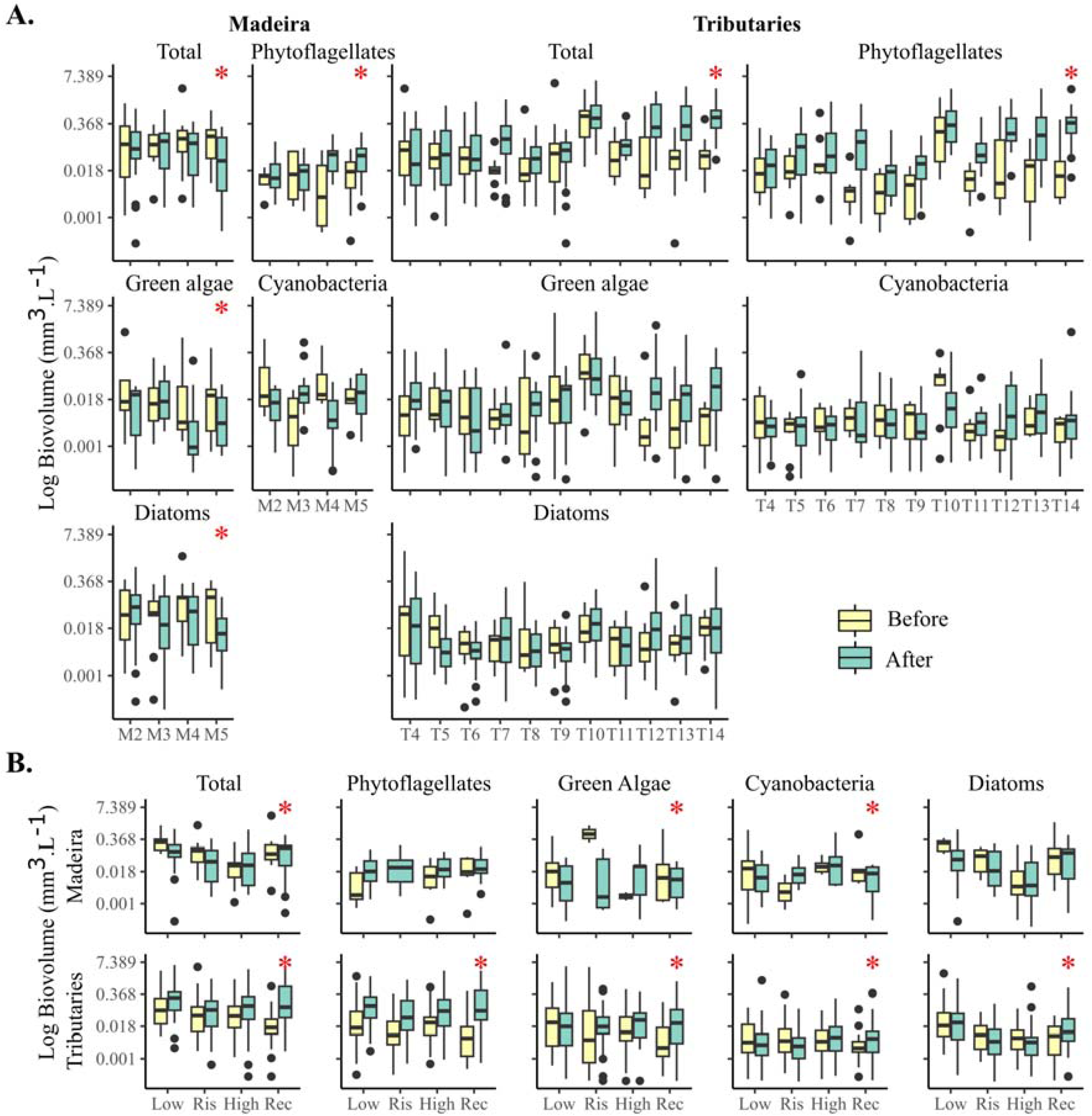
**A.** Difference in the distribution of total biomass and for each group in the tributaries and main channel during the construction phases of the dam. Asterisks represent significant differences. B. Differences in the biomass distribution of the total community and of each group during the hydrological periods and construction phases of the dam. Asterisks indicate significant differences in the biomass values obtained.

### 3.3 Relationship between abiotic variables and community biomass

The GLMM also revealed that the factors leading total biomass differed between the main channel and its tributaries. (Figure 6 Table S5). Moreover, the significance of these factors varied across taxonomic groups, depending on whether the occurrence or the biomass data were considered (Figure 6).

**Figure 6:**
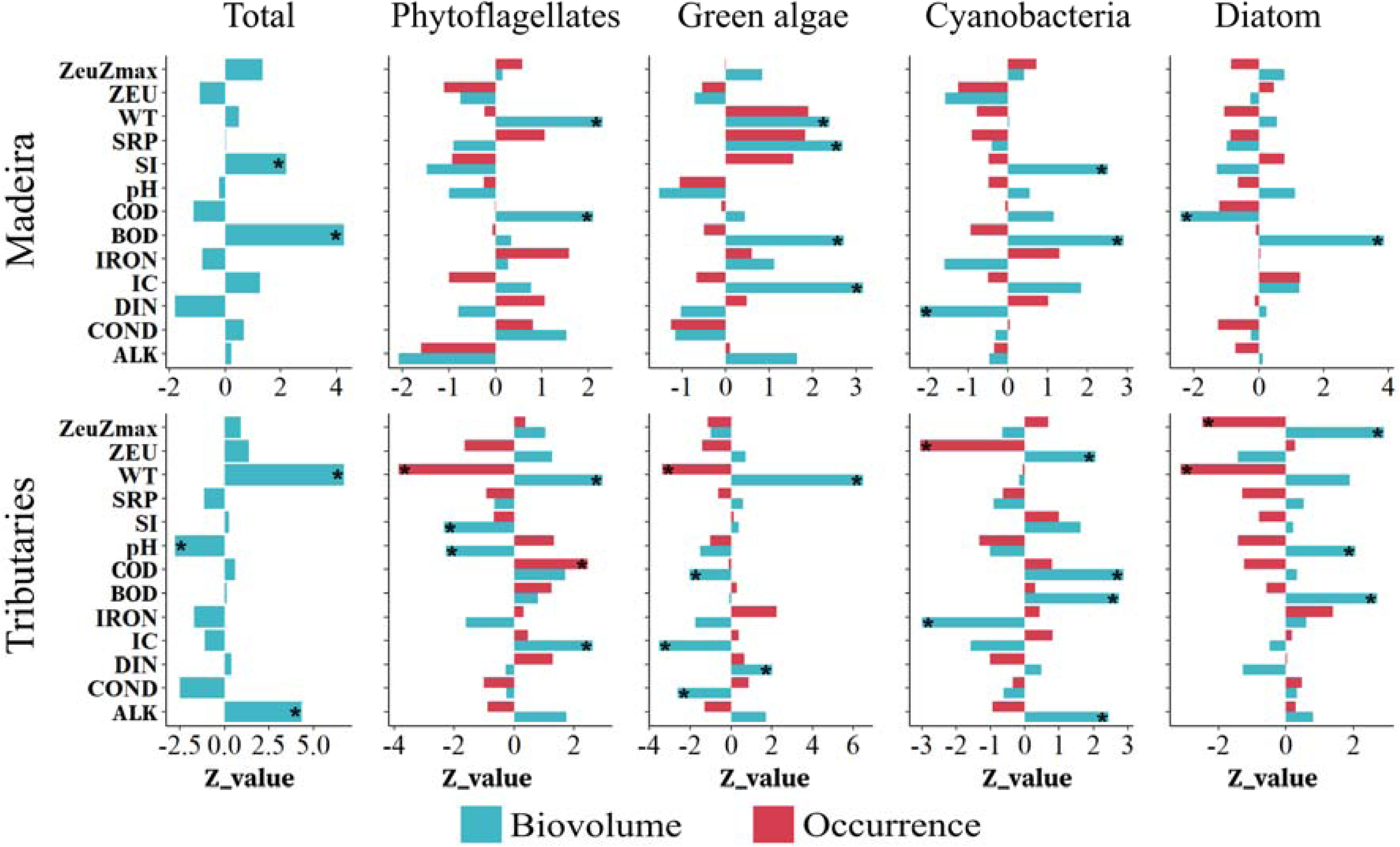
Influence of abiotic variables on total biomass and on the biomass and occurrence of algae groups from the main channel and tributaries. Asterisks represent a significant effect. Biomass is shown by blue bars while occurrence is shown by red bars. Zeu:Zmax – light availability in the water column; Zeu – euphotic zone; WT - water temperature; SRP – soluble reactive phosphorous; SI - Silica; PH - hydrogen potential; COD – chemical oxygen demand; BOD – biochemical oxygen demand; IC – dissolved inorganic carbon; DIN – dissolved inorganic nitrogen; COND – Conductivity; Alk – alkalinity.

## 4 Discussion

Several dams will occupy rivers in the Amazon region in the coming decades. Understanding their environmental implications and the subsequent effect on organisms is crucial for developing effective management and conservation strategies, as for anticipating and mitigating potential future. In this study, we demonstrated that the Jirau dam, as expected, induced changes in the biomass and structure of the phytoplankton communities in both the main channel and tributaries, corroborating our hypotheses. Specifically, damming promoted increased phytoplankton biomass in the tributaries, driven by the proliferation of species adapted to lacustrine, particularly phytoflagellates. Our findings highlight that while hydroelectric projects with Run-of-River (ROR) dams have historically been associated with minimal environmental impact, they can still significantly alter biotic characteristics, even in main channels where ROR dams are presumed to have less impact. This section delves into the impacts of damming on phytoplankton community dynamics and total biomass. We also discuss broader implications, including the potential future degradation of the Amazonian region, and how our results contribute to preventing undesirable scenarios.

### 4.1 Changes in phytoplankton structure

The construction of the Jirau dam seemed to influence changes in the phytoplankton communities in both the main channel and its tributaries, increasing the variation in communities at all hydrological seasons. This contrasts with previous research in the same study zone, which showed that biotic variation of planktonic communities decreased after damming (Pineda et al., 2024; Zanon et al., 2024). Unlike that research, our analysis focused on community variation between damming phases, considering each hydrological season separately. Figure 4 shows both spatial (between sampled points) and temporal (samples from different periods but within the same hydrological season and damming phase) variations together. However, given that the temporal water level variation decreased after damming and that spatial variation increased due to the emergence of lentic regions (which contrast with the lotic regions), the observed increase in sample dispersion was mainly related to spatial variation rather than temporal variation within the same hydrological season. Our results, together with research findings of Pineda et al. (2024) and Zanon et al. (2024), suggest that after damming, there could be an increase in spatial variation while temporal variation decreases. However, the spatial variation does not seem to be enough to compensate for the loss of temporal variation since the general pattern exhibit reduction in biotic heterogeneity (Pineda et al., 2024).

The increase in spatial variation that we observed can be attributed to the mosaic of habitats with varying features that resulted after damming. Following damming, as in our case, a region impacted by run-of-river projects shows not just rivers but also habitats with lake-like properties because of backwaters formed at the tributary’s junction with the main channel (Almeida et al., 2019). Each one of those habitats can then favor different species with different adaptations (Reynolds et al., 2002; Rodrigues et al., 2018; Zanco et al., 2017) and increase the spatial variation in communities. Our results indicated that after damming the number of indicator species of tributaries increased by 300%. It means that the characteristics of tributaries favored different species than those favored by the main channel. Furthermore, most of the species identified as tributary indicators after damming are typical of standing waters (Supplementary material), whereas most indicator species at the main channel were associated with lotic environments. It is consistent with our evidence (Supplementary material – Figure S1) and from predictions from other studies (e.g., Fearnside, 2013) about the stratification of tributaries after damming.

### 4.2 Changes in phytoplankton biomass

In the case of the total biomass, it decreased in the main channel (which was unexpected) and increased in the tributaries (as expected). We believe that the reduction of biomass in the main channel was caused by a decrease in phytoplankton export from tributaries, as the dam in the Madeira River increased the water level and reduced the water flow in the tributaries. However, more research is necessary to study the effect of damming on the dispersion between tributaries and the main channel. In any case, our result contrasts with evidence showing that run-of-river projects have little effect on large rivers (Almeida et al., 2019), and suggests that this kind of project may affect the flux of matter and energy in dammed rivers by decreasing the biomass of primary producers in the main channel.

We attributed the biomass increase in the tributaries to a reduction in their water flow, which increase water residence time, promotes stratification, and consequently favors phytoplankton development. As support for that assumption, we demonstrated an increase in species associated with standing waters as well as evidence of water stratification at certain tributaries after damming (Supplementary material – Figure S1). Additionally, we observed that water temperature had a positive effect on total biomass, and that temperature increased after damming, which may relate to stratification. Moreover, the stratification in tributaries in Madeira River has been documented after the construction of the Santo Antônio hydropower project (Almeida et al., 2019), which is also a large ROR located downstream of the Jirau dam. Unfortunately, we did not have stratification data for each sampling to include in our biomass models. Nevertheless, our observations suggest that even a run-of-river dam can lead to changes in the hydrodynamics of affected basins, mainly at tributaries, and that these changes can increase the phytoplankton biomass in tributaries.

Although total biomass increased at the tributaries, not all the phytoplankters reacted to damming in the same way. While the biomass of flagellates increased, that of cyanobacteria decreased. We believe that while lentic characteristics appeared to favor the whole phytoplankton community, chemical factors appeared to determine which group might thrive best. For instance, we found that an increase in nitrogen concentration following damming had a negative effect on cyanobacteria growth but favored green algae. In another case, high phosphate concentrations boosted cyanobacteria while harming phytoflagellates. In both examples, while the positive effect was associated with increased biomass, the negative effect may be linked to the rise in biomass of antagonistic groups, which dominate and limit the growth of other phytoplankton groups (Reynolds, 2006).

Phytoflagellates dominated the biomass, which may be related to the accumulation of organic material in the backwater regions of the tributaries. Despite the vegetation removal practices implemented before the reservoir filling in the Jirau Project (Energia Sustentável do Brasil, 2020), it seems that the flooded area still covered a significant amount of vegetation biomass (Cochrane et al., 2017), affecting the plant community and the decomposition of organic matter (Medeiros et al., 2023). The 10-meter average increase in the water level of the Madeira River submerged more than 160 km² of natural forest (Cochrane et al., 2017). We observed that chemical oxygen demand (COD) increased after damming in tributaries, indicating that the process of degradation of organic matter consumed more oxygen with damming. In this way, organisms with the ability of take advantage of a high amount of organic matter could domain. As changes in water features influence all aquatic species, these alterations can have an impact on the entire trophic web. For instance, a decrease in oxygen can relate to the death of fish (Demertzioglou et al., 2022) and the favoring of invasive species more tolerant to those new environmental conditions, such as some macrophyte species (Harrow-Lyle and Kirkwood, 2021).

The increase in algal biomass that we have identified here may not be restricted to our study zone. A comparable rise in phytoplankton biomass in backwater was registered in the influence zone of the dam of the Santo Antônio dam (Appel, 2017), downstream Jirau dam. As in our case, damming caused the formation of standing waters in that location (Almeida et al., 2019), which likely supported algal development as in our case. Furthermore, an increase in phytoplankton biomass may be predicted through the amazon, since the Brazilian government plans to expand hydroelectric power generation in the Brazilian Amazon over the next 25 years (Nobre et al., 2016) with the construction of thirty large dams (Empresa de Pesquisa Energética, 2013) and the flooding of ∼12,000 km^2^ (Nobre et al., 2016).

### 4.3 What our findings mean for the future and how we can work to prevent

Our study revealed that the creation of backwaters post-damming in tributaries appeared to favor an increase in algal biomass, resembling lake-like conditions. Despite this increase, the biomass values remained low, indicative of oligotrophic to mesotrophic conditions (Vollenweider, 1968). This limited biomass was primarily limited by low phosphorus availability (less than 0.001µg.L^-1^). Our findings suggest that while standing water conditions created by the Jirau dam enhanced hydrodynamic conditions for phytoplankton growth, current nutrient levels are insufficient to support high and problematic phytoplankton biomass, at least in the present nutrient conditions.

However, we caution that the optimal hydrodynamic conditions created by the dam, such as increased residence time and stratification in tributaries, could potentially synergize with future nutrient increases in Amazonian rivers. Studies (e.g., dos Santos et al., 2014) indicate that nutrient enrichment could lead to elevated phytoplankton biomass levels, like issues observed in other dam-affected regions worldwide (Grill et al., 2019; Winemiller et al., 2016).

Human activities associated with nutrient enrichment in water have reached areas of the Amazon that would have been unimaginable a few decades ago. For example, deforestation for pastures has been linked to phosphorus rise in rivers like the Ji-Paraná River in the Amazon basin (Ballester et al., 2003). Furthermore, projections suggest that by 2050, pastures and agricultural expansion could cover 40% of the entire Amazon region (Soares-Filho et al., 2006), potentially increasing nutrient loads to waterways. Urban expansion facilitated by dams and agro-industrial operations has also contributed to nutrient increases in Amazonian rivers due to untreated sewage (Ferreira et al., 2021).

Given these observations, we urge authorities to implement stringent protocols for nutrient management in future agricultural and urban projects in dam-influenced zones of the Amazon. Proper management practices can mitigate the risks associated with phytoplankton blooms, preserving water quality and the ecosystem services derived from aquatic environments, including water supply, recreation, and fishing. This proactive approach to nutrient management is essential to safeguarding the ecological integrity of Amazonian rivers.

## 5 Conclusion

Even run-of-the-river hydroelectric projects like the Jirau hydroelectric plant can alter water chemistry and hydrodynamics, subsequently affecting the characteristics of microbial communities such structure and biomass of phytoplankton. This impact could potentially extend across the Amazon region, as several similar hydropower projects are currently under construction in the area. Given that the flooding of large swathes of vegetation is a common consequence of these projects, it is anticipated that dams in the Amazon region will lead to an increase in mixotrophic organisms such as flagellates, which take advantage of organic matter. Additionally, the dams are expected to elevate the chemical demand for oxygen to degrade the accumulated organic matter, thereby reducing oxygen availability. While damming does promote the biomass of phytoplankton due to extended residence times and water column stratification, it may not suffice to produce problematic levels of biomass due to limited nutrient availability. Therefore, authorities must remain vigilant regarding activities that may exacerbate these issues and impact water quality in the Amazon. This includes monitoring practices that could result in heightened nutrient loading in aquatic habitats, such as agricultural practices, urban development, industrial activities, and sewage discharge.

## Acknowledgments

We thank the Programa de Pós-graduação em Ecologia de Ambientes Aquáticos Continentais and the Núcleo de Pesquisas em Limnologia, Ictiologia e Aquicultura (Nupélia) from the Universidade Estadual de Maringá (UEM) for supplying infrastructure. We thank the Coordenação de Aperfeiçoamento de Pessoal de Nível Superior— Brazil (CAPES) for the scholarship granted to BMC, FMZ and RFS. AP thanks Conselho Nacional de Desenvolvimento Científico e Tecnológico (CNPq) for postdoctoral fellowships. LCR was supported by productivity fellowships from CNPq. The authors state that they have no conflicts of interest relating to this study.

## Declarations

### Data availability

The datasets generated during and/or analyzed during the current study are available from the corresponding author on reasonable request

### Consent to participate

Not applicable.

### Consent to publication

Not applicable

### Ethical approval

Not applicable. The present study did not involve research with humans or animals

---

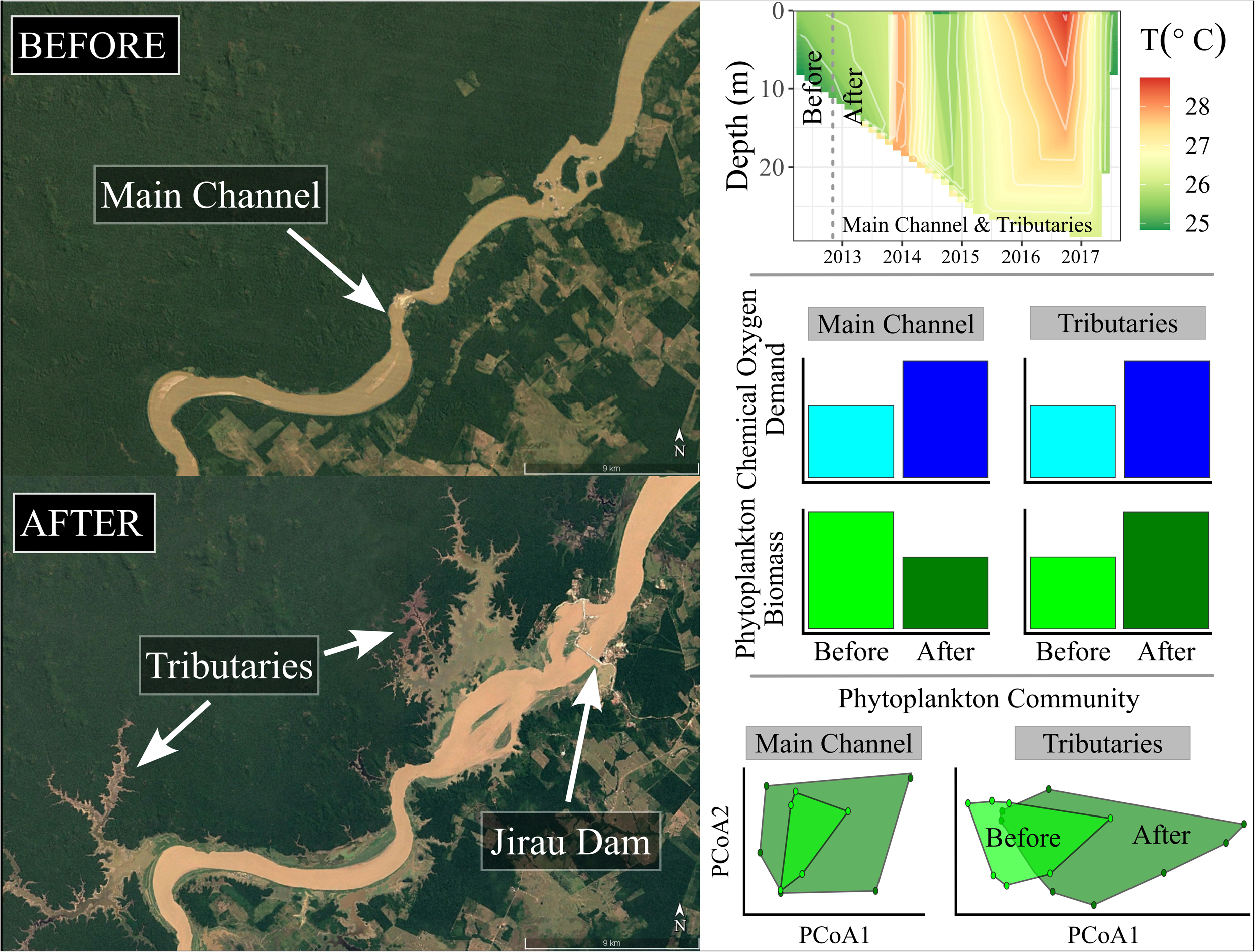

