## Supplementary material for "Impact of Run-of-River Damming on Increasing Phytoplankton Biomass and Species Shift in a Large Amazonian River" (from another version of this preprint: version 2, 2024-06-28)

**Figure S1 –** Fluctuations in the water level of the Madeira River (reservoir) from 2008 to 2016, during the construction phases of the Jirau dam. The boxplots depict the distribution of the data, with the box representing the interquartile range, the horizontal bar indicating the median, and the vertical bars showing the minimum and maximum values, with any outliers represented as individual points. It is noteworthy that following the completion of the dam, the water level not only rose, but also experienced a decrease in variation.

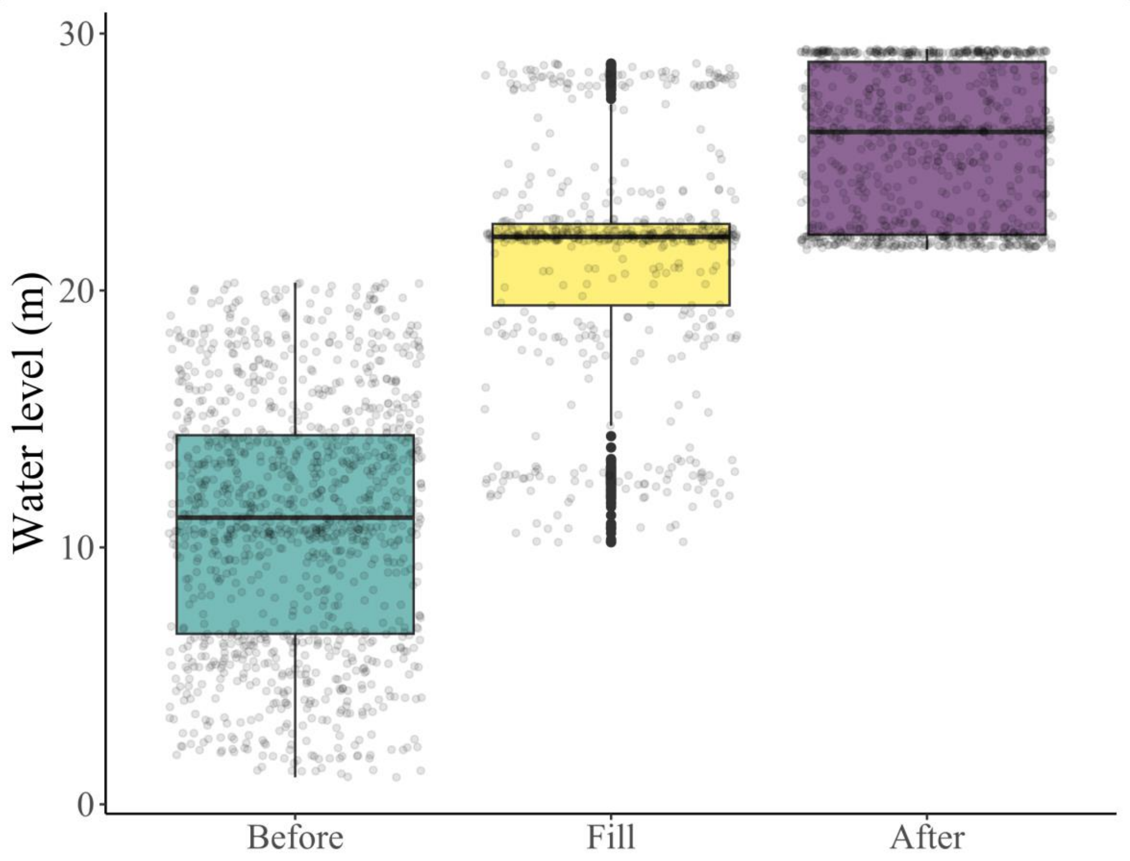

**Figure S2 -** Graphical representation of thermal stratification for some points of tributaries and Madeira River.

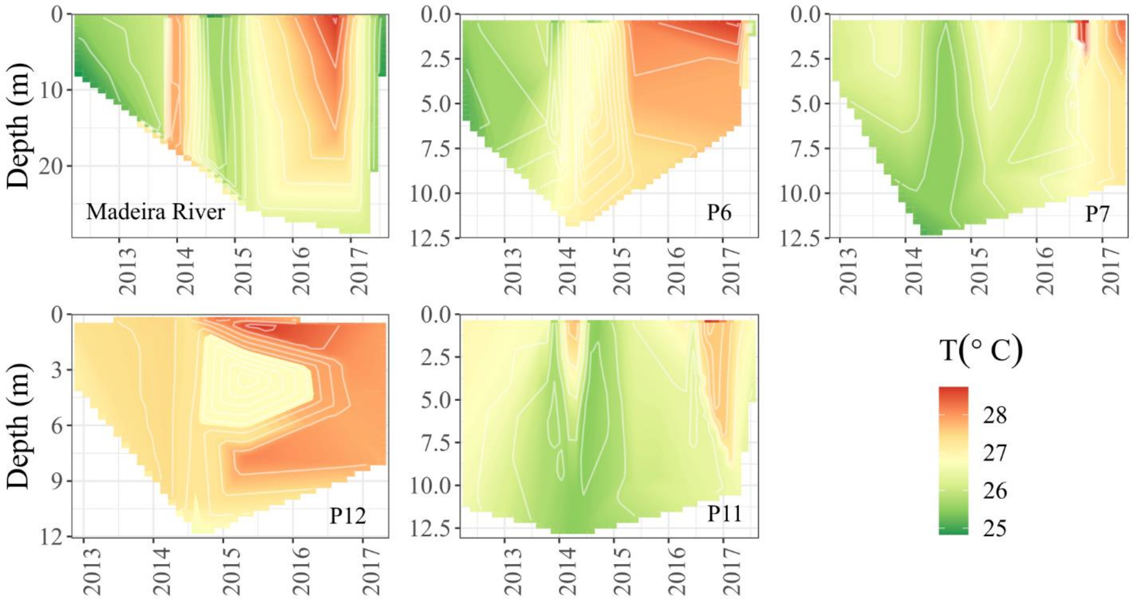


**Table S1**. Effect of damming phases on the environmental factors at Madeira River and tributaries. Before damming was considered as a reference. A negative and significant Estimate means that values of the variable increase after damming. Bold highlights significant effects of damming phases.

| | Madeira River | | | Tributaries | | |
|---|---|---|---|---|---|---|
| **Variable** | **Estimate** | **z value** | **P** | **Estimate** | **z value** | **P** |
| PO4 | -4.2670 | -1.37 | 1.70E-01 | 2.3280 | 0.32 | 7.47E-01 |
| TW | 0.0004 | 0.50 | 6.29E-01 | **-0.0414** | **-5.70** | **1.41E-08** |
| DIN | 0.0334 | 0.51 | 6.08E-01 | -0.0763 | -1.81 | 6.96E-02 |
| Depth | -0.1352 | -1.77 | 7.68E-02 | **-0.1813** | **-4.87** | **1.12E-06** |
| Conductivity | **0.4558** | **3.28** | **1.05E-03** | **-0.5927** | **-8.44** | **<2E-16** |
| COD | **-0.3312** | **-2.60** | **9.21E-03** | **-0.3088** | **-4.60** | **4.14E-06** |
| BOD | -0.0274 | -0.47 | 6.36E-01 | -0.0131 | -0.23 | 8.22E-01 |
| Zeu | **2.4751** | **4.32** | **1.55E-05** | **0.1820** | **4.20** | **2.66E-05** |
| Iron | **0.8246** | **6.09** | **1.14E-09** | **0.1079** | **3.89** | **9.92E-05** |
| Alkalinity | **0.2559** | **2.89** | **3.91E-03** | **-0.3157** | **-6.02** | **1.79E-09** |
| IC | **0.1366** | **2.19** | **2.84E-02** | **-0.4285** | **-11.05** | **<2E-16** |
| Silica | **0.5097** | **3.19** | **1.41E-03** | -0.0547 | -1.44 | 1.50E-01 |
| pH | **0.5877** | **5.10** | **3.46E-07** | 0.1177 | 1.47 | 1.41E-01 |
| Zeu:Zmax | **16.0230** | **2.76** | **5.81E-03** | **0.8814** | **6.58** | **4.77E-11** |

**Table S2:** GLMM result of the interaction between phases:hydrological period for all groups in the tributaries and main channel.

**Main channel**

| | **Greens** | | | **Cyanobacteria** | | | **Total biomass** | | |
|---|---|---|---|---|---|---|---|---|---|
| | **F** | **Df** | **Pr(>F)** | **F** | **Df** | **Pr(>F)** | **F** | **Df** | **Pr(>F)** |
| Hydrological period | 6.49 | 3.00 | 0.09 | 11.38 | 3.00 | **<0.001** | 13.45 | 3.00 | **<0.001** |
| Phases | 12.69 | 1.00 | **<0.001** | 2.03 | 1.00 | 0.15 | 6.00 | 1.00 | **0.01** |
| Hydrological period:phases | 17.29 | 3.00 | **<0.001** | 18.07 | 3.00 | **<0.001** | 8.15 | 3.00 | **0.04** |

**Tributaries**

| | **Greens** | | | **Phytoflagellates** | | | **Diatoms** | | | **Cyanobacteria** | | | **Total biomass** | | |
|---|---|---|---|---|---|---|---|---|---|---|---|---|---|---|---|
| | **F** | **Df** | **Pr(>F)** | **F** | **Df** | **Pr(>F)** | **F** | **Df** | **Pr(>F)** | **F** | **Df** | **Pr(>F)** | **F** | **Df** | **Pr(>F)** |
| Hydrological period | 29.40 | 3.00 | **<0.001** | 24.54 | 3.00 | **0.00** | 73.29 | 3.00 | **<0.001** | 12.45 | 3.00 | **<0.001** | 26.58 | 3.00 | **<0.001** |
| Phases | 1.35 | 1.00 | 0.25 | 79.70 | 1.00 | **<0.001** | 0.02 | 1.00 | 0.89 | 5.30 | 1.00 | **0.02** | 24.38 | 1.00 | **<0.001** |
| Hydrological period:phases | 12.79 | 3.00 | **0.01** | 19.68 | 3.00 | **<0.001** | 10.96 | 3.00 | **0.01** | 23.87 | 3.00 | **<0.001** | 56.01 | 3.00 | **<0.001** |

**Table S3:** GLMM result of the hydrological period and phases for all groups in the tributaries and main channel.

| Main channel | | | | | | | | | | | | | | | |
|---|---|---|---|---|---|---|---|---|---|---|---|---|---|---|---|
| | **Greens** | | | **Phytoflagellates** | | | **Diatoms** | | | **Cyanobacteria** | | | **Total biomass** | | |
| | **F** | **Df** | **Pr(>F)** | **F** | **Df** | **Pr(>F)** | **F** | **Df** | **Pr(>F)** | **F** | **Df** | **Pr(>F)** | **F** | **Df** | **Pr(>F)** |
| Phases | 18.94 | 1.00 | **<0.001** | 5.34 | 1.00 | **0.02** | 4.99 | 1.00 | **0.03** | 1.51 | 1.00 | 0.22 | 9.63 | 1.00 | **<0.001** |
| Hydrological period | 8.40 | 3.00 | **0.04** | 3.57 | 3.00 | 0.31 | 26.49 | 3.00 | **<0.001** | 10.37 | 3.00 | **0.02** | 13.18 | 3.00 | **<0.001** |

| Tributaries | | | | | | | | | | | | | | | |
|---|---|---|---|---|---|---|---|---|---|---|---|---|---|---|---|
| | **Greens** | | | **Phytoflagellates** | | | **Diatoms** | | | **Cyanobacteria** | | | **Total biomass** | | |
| | **F** | **Df** | **Pr(>F)** | **F** | **Df** | **Pr(>F)** | **F** | **Df** | **Pr(>F)** | **F** | **Df** | **Pr(>F)** | **F** | **Df** | **Pr(>F)** |
| Phases | 0.03 | 1.00 | 0.87 | 58.25 | 1.00 | **<0.001** | 0.53 | 1.00 | 0.47 | 3.24 | 1.00 | 0.07 | 7.35 | 1.00 | **0.01** |
| Hydrological period | 23.33 | 3.00 | **<0.001** | 20.18 | 3.00 | **<0.001** | 73.65 | 3.00 | **<0.001** | 9.89 | 3.00 | **0.02** | 15.84 | 3.00 | **<0.001** |

**Table S4:** GLMM result of the sum between phases and hydrological period for diatoms in Madeira River.

| | **F** | **Df** | **Pr(>F)** |
|---|---|---|---|
| Hydrological period | 24.71 | 3.00 | **<0.001** |
| Phases | 3.60 | 1.00 | 0.06 |
| Hydrological period+phases | 0.99 | 3.00 | 0.80 |

**Table S5:** GLMM result of environmental variables by phytoplankton groups and biomass total. (Zeu:Zmax, pH, COND, Zeu, SRP, DIN, BOD, COD, Iron, Alk, IC, WT, Si).

**Main channel**

| | **Total biomass** | | | **Phytoflagellates** | | | **Greens** | | | **Cyanobacteria** | | | **Diatoms** | | |
|---|---|---|---|---|---|---|---|---|---|---|---|---|---|---|---|
| | **Estimate** | **Z** | **Pr(>\|z\|)** | **Estimate** | **Z** | **Pr(>\|z\|)** | **Estimate** | **Z** | **Pr(>\|z\|)** | **Estimate** | **Z** | **Pr(>\|z\|)** | **Estimate** | **Z** | **Pr(>\|z\|)** |
| $Z_{eu}$:$Z_{max}$ | 1.08 | 1.33 | 0.18 | 0.12 | 0.16 | 0.88 | 1.29 | 0.84 | 0.40 | 0.64 | 0.41 | 0.68 | 0.73 | 0.80 | 0.43 |
| pH | -0.03 | -0.22 | 0.83 | -0.20 | -1.00 | 0.32 | -0.45 | -1.53 | 0.13 | 0.16 | 0.54 | 0.59 | 0.22 | 1.11 | 0.27 |
| COND | 0.16 | 0.68 | 0.50 | 0.56 | 1.53 | 0.13 | -0.55 | -1.16 | 0.25 | -0.16 | -0.31 | 0.76 | -0.06 | -0.24 | 0.81 |
| Zeu | -0.45 | -0.91 | 0.36 | -0.28 | -0.76 | 0.45 | -0.73 | -0.72 | 0.47 | -1.35 | -1.57 | 0.12 | -0.15 | -0.26 | 0.80 |
| SRP | 0.00 | 0.03 | 0.98 | -0.20 | -1.49 | 0.14 | 0.00 | -0.01 | 0.99 | 0.43 | 2.52 | **0.01** | -0.14 | -1.30 | 0.19 |
| DIN | -0.17 | -1.80 | 0.07 | -0.14 | -0.79 | 0.43 | -0.31 | -1.03 | 0.30 | -0.39 | -2.21 | **0.03** | 0.02 | 0.24 | 0.81 |
| BOD | 0.52 | 4.25 | **<0.001** | 0.07 | 0.35 | 0.73 | 0.64 | 2.72 | **0.01** | 0.52 | 2.91 | **0.00** | 0.51 | 3.86 | **<0.001** |
| COD | -0.16 | -1.13 | 0.26 | 0.45 | 2.11 | **0.04** | 0.16 | 0.44 | 0.66 | 0.40 | 1.16 | 0.25 | -0.38 | -2.42 | **0.02** |
| Iron | -0.07 | -0.82 | 0.41 | 0.06 | 0.27 | 0.79 | 0.32 | 1.12 | 0.26 | -0.44 | -1.60 | 0.11 | 0.00 | -0.02 | 0.99 |
| Alk | 0.04 | 0.22 | 0.82 | -0.66 | -2.08 | **0.04** | 0.60 | 1.65 | 0.10 | -0.15 | -0.46 | 0.64 | 0.02 | 0.11 | 0.91 |
| IC | 0.14 | 1.25 | 0.21 | 0.14 | 0.77 | 0.44 | 0.65 | 3.16 | **<0.001** | 0.39 | 1.85 | 0.06 | 0.17 | 1.26 | 0.21 |
| WT | 0.07 | 0.50 | 0.62 | 0.44 | 2.30 | **0.02** | 0.73 | 2.39 | **0.02** | 0.01 | 0.04 | 0.97 | 0.11 | 0.56 | 0.58 |
| Si | 0.15 | 2.18 | **0.03** | -0.16 | -0.90 | 0.37 | 0.48 | 2.69 | **0.01** | -0.07 | -0.40 | 0.69 | -0.09 | -1.00 | 0.32 |

**Tributaries**

| | **Total biomass** | | | **Phytoflagellates** | | | **Greens** | | | **Cyanobacteria** | | | **Diatoms** | | |
|---|---|---|---|---|---|---|---|---|---|---|---|---|---|---|---|
| | **Estimate** | **Z** | **Pr(>\|z\|)** | **Estimate** | **Z** | **Pr(>\|z\|)** | **Estimate** | **Z** | **Pr(>\|z\|)** | **Estimate** | **Z** | **Pr(>\|z\|)** | **Estimate** | **Z** | **Pr(>\|z\|)** |
| $Z_{eu}$:$Z_{max}$ | 0.09 | 0.93 | 0.35 | 0.10 | 1.05 | 0.30 | -0.14 | -1.01 | 0.31 | -0.08 | -0.65 | 0.52 | 0.30 | 2.90 | **<0.001** |
| pH | -0.21 | -2.77 | **0.01** | -0.19 | -2.28 | **0.02** | -0.19 | -1.52 | 0.13 | -0.12 | -1.03 | 0.31 | 0.18 | 2.06 | **0.04** |

| | | | | | | | | | | | | | | | |
|---|---|---|---|---|---|---|---|---|---|---|---|---|---|---|---|
| COND | -0.32 | -2.51 | **0.01** | -0.05 | -0.27 | 0.79 | -0.75 | -2.62 | **0.01** | -0.14 | -0.63 | 0.53 | 0.09 | 0.32 | 0.75 |
| Zeu | 0.15 | 1.36 | 0.17 | 0.14 | 1.27 | 0.20 | 0.11 | 0.70 | 0.48 | 0.29 | 2.06 | **0.04** | -0.16 | -1.42 | 0.16 |
| SRP | -0.15 | -1.12 | 0.26 | -0.34 | -2.34 | **0.02** | 0.06 | 0.37 | 0.71 | 0.47 | 1.63 | 0.10 | 0.05 | 0.21 | 0.83 |
| DIN | 0.05 | 0.39 | 0.70 | -0.04 | -0.29 | 0.77 | 0.43 | 2.02 | **0.04** | 0.08 | 0.48 | 0.63 | -0.18 | -1.28 | 0.20 |
| BOD | 0.01 | 0.12 | 0.91 | 0.07 | 0.80 | 0.43 | -0.01 | -0.08 | 0.94 | 0.22 | 2.75 | **0.01** | 0.22 | 2.70 | **0.01** |
| COD | 0.05 | 0.61 | 0.55 | 0.13 | 1.71 | 0.09 | -0.27 | -2.03 | **0.04** | 0.33 | 2.87 | **<0.001** | 0.03 | 0.32 | 0.75 |
| Iron | -0.38 | -1.70 | 0.09 | -0.45 | -1.61 | 0.11 | -0.88 | -1.76 | 0.08 | -0.87 | -2.99 | **<0.001** | 0.17 | 0.59 | 0.55 |
| Alk | 0.70 | 4.35 | **<0.001** | 0.34 | 1.75 | 0.08 | 0.41 | 1.71 | 0.09 | 0.57 | 2.44 | **0.01** | 0.18 | 0.81 | 0.42 |
| IC | -0.11 | -1.10 | 0.27 | 0.31 | 2.61 | **0.01** | -0.48 | -3.51 | **<0.001** | -0.20 | -1.58 | 0.12 | -0.06 | -0.49 | 0.62 |
| WT | 0.53 | 6.68 | **<0.001** | 0.27 | 2.95 | **<0.001** | 0.83 | 6.47 | **<0.001** | -0.02 | -0.17 | 0.87 | 0.16 | 1.88 | 0.06 |
| Si | 0.05 | 0.24 | 0.81 | -0.12 | -0.66 | 0.51 | 0.21 | 0.60 | 0.55 | -0.24 | -0.91 | 0.36 | 0.12 | 0.53 | 0.60 |
